# Structure and Dynamics of the HIV-1 Envelope Protein on the Virion Envelope

**DOI:** 10.64898/2026.05.18.725998

**Authors:** Ayan Majumder, Mandira Dutta, Levi Cherek, Gregory A. Voth

## Abstract

HIV-1 buds from infected cells as immature virion particles with a scattered envelope glycoprotein (Env) distribution on their envelope. It then undergoes maturation, during which the viral protease cleaves the Gag polyprotein at multiple sites, leading to structural reorganization of the viral particle and lateral redistribution of Env proteins, ultimately rendering the virion infectious. However, the underlying mechanism of maturation-induced Env reorganization remains elusive. In this study, we combine microsecond-long all-atom (AA), bottom-up coarse-grained (CG) molecular dynamics simulations, and diffusion model-based backmapping to investigate the structural organization and key interactions of Env in viral membranes. AA simulations of fully glycosylated Env embedded in HIV-1 mimetic asymmetric bilayers were first performed to characterize its conformational dynamics and Env-lipid interactions. We then developed a bottom-up CG model of glycosylated Env from that AA data and simulated the mature HIV-1 virion envelope containing multiple Env proteins. The CG simulations predict that Env proteins form clusters through interactions mediated by the cytoplasmic tail domain (CTD) and adopt diverse tilted conformations within these clusters. These CG simulations were then backmapped to AA resolution and further AA simulations were carried out to identify, in detail, the specific interacting residues in the Env clusters. Additionally, analysis of epitope accessibility shows that broadly neutralizing antibodies (bnAbs) targeting the V1/V2 and V3 loops may efficiently interact with Env clusters on the mature virion surface. Together, these results provide a molecular mechanism for Env oligomerization during viral maturation and offer new insights into the accessibility of bnAb epitopes on Env clusters.

## Introduction

The human immune deficiency virus-1 (HIV-1) lipid envelope carries the viral envelope protein (Env), which mediates viral entry into the host cell and initiates infection.^1, 2^ Env on the HIV-1 surface is the sole target of neutralizing antibodies and an important target for vaccine design efforts.^3-5^ Env is initially translated as gp160, a precursor protein containing approximately 850 residues. The gp160 protein trimerizes, undergoes glycosylation, and is subsequently cleaved by a furin protease into gp120 and gp41 subunits, which mediate host cell receptor binding and membrane fusion, respectively.^6^ Gp120 and gp41 remain as a noncovalently associated heterodimer, with gp41 anchored in the viral membrane through interactions mediated by the membrane-proximal external region (MPER), transmembrane domain (TMD), and cytoplasmic tail (CT).^7^ The heavily glycosylated trimer of gp120-gp41 heterodimer forms a functional spike on the viral surface.^8, 9^ HIV-1 entry into the host cell begins with gp120 binding to the host cell receptor CD4 and the coreceptors CCR5 or CXCR4.^1, 10^ This interaction triggers a cascade of structural reorganization in gp120 and gp41, ultimately leading to the insertion of the gp41 fusion peptide into the host cell and then fusion of the viral and host cell membranes.^11, 12^

Env is incorporated into the nascent virion membrane during viral assembly at the plasma membrane. A typical HIV-1 virion exhibits a relatively low number of spike proteins compared to other enveloped viruses, with only 7-14 spikes per virion.^13, 14^ HIV-1 assembly at the plasma membrane begins with the binding of Gag polyprotein to the inner-leaflet of the plasma membrane mediated by the interactions between the matrix (MA) domain and phosphatidylinositol 4,5-biphosphate (PIP2) lipids.^15^ Gag polyprotein then oligomerizes to form a viral assembly lattice. Experimental studies using fluorescence microscopy have shown that Env proteins are specifically recruited to the vicinity of the assembly site, and deletion of the Env CT abrogates Env incorporation.^16^ At the final stage of viral assembly, Gag recruits the endosomal sorting complex required for transport (ESCRT) machinery, leading to budding and the release of immature virions.^17^ Concurrently or shortly after budding from the plasma membrane, the Gag lattice on the cytofacial side of the viral membrane disassembles into its constituent domains via a tightly regulated protease-mediated cleavage pathway. This process results in substantial structural rearrangements of the viral particle, termed viral maturation.^18, 19^ Maturation-induced lateral reorganization of Env has been proposed to render viral infectivity;^20-22^ however, the underlying mechanism of Env clustering in mature virions remains elusive. Consistent with this notion, an immature virion, despite possessing the same number of Env proteins as a mature virion and with no apparent difference in Env structure, displays reduced efficacy in entering a target cell.^20, 21^ This effect could be caused by the stiffness of the immature Gag lattice underneath the viral envelope or by the reduced mobility of Env in immature virions.^23^ Kräusslich and coworkers have studied Env lateral organization using fluorescence microscopy and showed that maturation-induced coalescence of multiple Env proteins into a single cluster results in the formation of fully mature, infectious virions.^20^

A holistic model of the Env protein embedded in its viral membrane can provide insight into unresolved questions regarding Env dynamics at different stages of the HIV-1 lifecycle. Molecular dynamics (MD) simulations have been widely used to investigate protein dynamics in complex lipid bilayers.^24-28^ However, all-atom (AA) simulations of the full HIV-1 viral envelope, including HIV-1 assembly and budding at the plasma membrane remain largely inaccessible due to the high computational cost of simulating hundreds of millions of atoms over meaningful timescales. Bottom-up coarse-grained (CG) models^29^ have been employed to access the relevant timescales necessary to study large biomolecular assemblies beyond the reach of AA simulations (see examples in refs 30, 31). A systematic bottom-up CG model is generally constructed following a statistical mechanical framework to reproduce certain underlying thermodynamic properties of the corresponding AA simulations (e.g., the many-dimensional potential of mean force of the CG coordinates).^29^ We have previously developed a bottom-up CG model to simulate an asymmetric lipid bilayer mimicking the HIV-1 membrane composition.^32^ This CG model was shown to reproduce the properties of the AA asymmetric bilayer while being several orders of magnitude computationally faster than AA simulations.

In this study, we constructed a bottom-up CG model of the glycosylated Env protein embedded in an HIV-1 mimetic asymmetric lipid bilayer, based on data obtained from extensive AA MD simulations. Comparison of the CG model with the AA simulations shows that the CG model accurately reproduces the structural dynamics of Env and captures important Env-lipid interactions observed at the atomistic level. We then simulated a full CG HIV-1 virion envelope containing multiple copies of the Env protein to observe their clustering on the envelope. Additionally, we also simulated multiple copies of Env in HIV-1 mimetic flat asymmetric bilayers using our CG model. The results obtained from the simulations provide insight into the CT domain-mediated interactions governing Env clustering and the lateral organization of Env proteins in the mature virion, as well as the origins of the forces that drive these behaviors. We then backmapped the CG model to the AA level and through additional AA MD simulations of a backmapped Env dimer, we identified key interactions influencing the Env clustering. We also characterized the features that may influence the binding affinity of different neutralizing antibodies to Env clusters. Taken together, this detailed model of the Env on the full HIV-1 envelope establishes a framework for understanding largescale Env dynamics on the surface of the HIV-1 virion.

## Results and Discussion

### All-atom simulation of glycosylated Env in an asymmetric bilayer

We simulated a fully glycosylated Env embedded in HIV-1 virion-mimetic asymmetric bilayers. The full-length Env was constructed using the ectodomain structure resolved in the native virion envelope by Lee and coworkers^33^ using cryo-ET and the NMR structure of the membrane-interacting domain (MPER-TMD-CT) resolved by Chou and coworkers (**Figure 1**).^7^ A total of 72 distinct glycans were modeled on the Env ectodomain following previous experimental studies to generate the fully glycosylated full-length Env (**Figure 1 and Tables S1 and S2** of the Supporting Information).^34-37^ The HIV-1 mimetic asymmetric lipid bilayer was constructed following previous experimental and computational studies.^7, 32, 38, 39^ The exofacial leaflet was composed of 40 mol% LSM, 20 mol% POPC, and 40 mol% cholesterol, whereas the cytofacial leaflet consisted of 40 mol% POPE, 30 mol% POPS, 15 mol% PIP2, and 15 mol% cholesterol. A higher PIP2 concentration (∼7 mol%) than that reported in previous lipidomics experiments (∼3 mol%)^39^ was used to more thoroughly facilitate sampling protein-PIP2 interactions. Majumder et al.^32^ previously simulated an asymmetric bilayer of the same composition and showed that the exofacial leaflet is in a liquid-ordered state, while the cytofacial leaflet is a liquid-disordered state at room temperature, highlighting the cholesterol condensation effect. However, no lipid phase separation was observed in either leaflet.^32^

**Figure 1.**
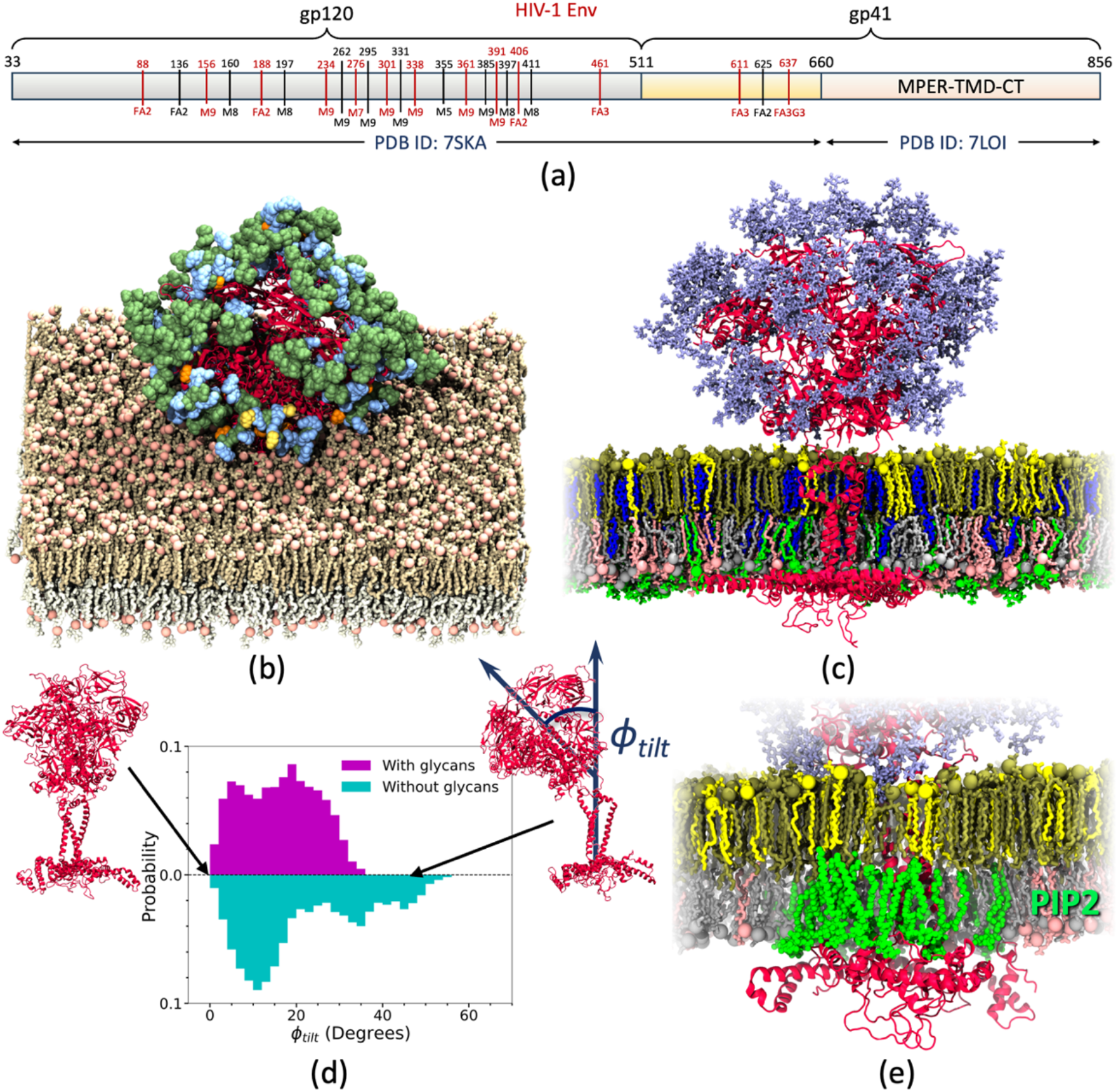
(a) Construct of the glycosylated Env protein simulated in this study. (b) Pictorial representation of glycans containing mannose (green), GlcNAc (blue), fucose (orange), and galactose (yellow) attached to the ectodomain of Env. (c) Side view of the glycosylated Env embedded in an asymmetric bilayer containing POPC (yellow), LSM (tan), and cholesterol (blue) in the exofacial leaflet, and POPE (gray), POPS (pink), PIP2 (green), and cholesterol in the cytofacial leaflet. The Env protein and glycans are represented in red and ice blue, respectively. (d) Tilt angle distribution of the Env ectodomain relative to the membrane normal, comparing systems with and without glycans. (e) Pictorial representation of PIP2 lipids (green) interacting with the CT domain of Env (red).

Previous computational studies of truncated and full-length Env have shown that Env anchors to the membrane with the MPER and CT domains of gp41 interacting with water at the membrane interface on the exofacial and cytofacial sides, respectively.^7, 40-42^ Following these studies, the initial configuration of the Env-embedded lipid bilayer was prepared by placing the MPER and CT domains at the membrane-water interfaces on the exofacial and cytofacial sides, respectively. Because lipids in the asymmetric bilayer do not undergo phase separation, all lipids were initially placed randomly around Env. The resulting protein-lipid system was simulated using the all-atom CHARMM36m force field.^43^ Three independent replicate simulations were performed for Env with and without glycans embedded in the asymmetric bilayer (**Table S3**). In all replicate simulations, the CT domain formed a stable baseplate at the interface between the cytofacial leaflet and water, whereas the MPER domain interacted with the exofacial leaflet-water interface, consistent with previous studies.^7, 44^

Prior experimental studies^41, 45^ using cryo-ET have shown that the Env ectodomain adopts tilted conformations with respect to the membrane normal in virus-like particles. Shehata et al. simulated the glycosylated Env protein and showed that glycans at positions 88 and 611 play a role in this tilting motion.^41^ Croft et al. have also simulated glycosylated Env and characterized Env tilting in mature and immature virions. However, they did not observe significant glycan-lipid interactions in their simulations that correlate with Env tilting.^45^

We calculated the tilt angle distribution of the Env ectodomain with respect to the membrane normal (**Figure 1d**). The results show a significant tilting motion of Env, consistent with previous studies.^41, 45^ We also calculated a contact map to characterize interactions between glycans and lipids observed in the simulations (**Figure S1**). The contact map shows interactions between glycans at positions 88, 611, and 625 with lipids in the exofacial leaflet. However, a similar tilting motion of the Env ectodomain was also observed in simulations of Env without glycans. These results suggest that tilting of the ectodomain is an intrinsic motion of Env that occurs both in the presence and absence of glycans, while N-glycans interact with the membrane as a consequence of ectodomain tilting. Nevertheless, glycan-lipid interactions may play a role in stabilizing tilted Env conformations relative to upright conformations.

Previous experimental studies using EPR and NMR have reported diverse conformations of the MPER and TMD, indicating substantial conformational flexibility in these regions.^7, 44, 46-49^ We previously studied two different protein constructs of the membrane-interacting domains of gp41, and using a machine-learning (ML) based approach we showed that the MPER-TMD adopts diverse structures consistent with previous various experimental structures.^40^ We calculated hinge angle distributions in full-length Env formed at the junction between the MPER and TMD (*ϕ*_*top*_), as well as near the C-terminal region of the TMD (*ϕ*_*bottom*_). The hinge angle distributions of MPER-TMD in glycosylated Env are shown in **Figure S2**. The results show similar *ϕ*_*bottom*_ distributions for both full-length and truncated membrane-interacting constructs of Env. However, the presence of the ectodomain attached to MPER in full-length Env restricts its bending motion, resulting in higher *ϕ*_*top*_values. Nevertheless, similar conformational diversity of MPER-TMD is observed in glycosylated Env as in truncated constructs, with MPER-TMD adopting both uninterrupted α-helical and helix-turn-helix structures.

### Bottom-up coarse-grained model reproduces important AA protein-lipid ensemble properties

Using all-atom (AA) trajectories as a reference, we constructed a bottom-up coarse-grained (CG) model of glycosylated Env embedded in a CG HIV-1 mimetic asymmetric bilayer (**Figure 2a and 2b**). POPC, POPE, POPS, and PIP2 lipids were mapped onto six CG beads, whereas cholesterol and LSM lipids were mapped onto three and seven CG beads, respectively^32^ (**Figure 2c**). The bonded force field parameters of the lipids were constructed using the multiscale coarse-graining (MS-CG) protocol,^29, 50-53^ and the non-bonded parameters were developed using both MS-CG and relative entropy minimization (REM) methods.^54^ The bottom-up CG lipid model was shown to accurately capture the area per lipid, the liquid-crystal (P_2_) order parameters of the lipid tails, and the lateral lipid density along the membrane normal, as observed in all-atom simulations of HIV-1 mimetic asymmetric bilayers.^32^

**Figure 2.**
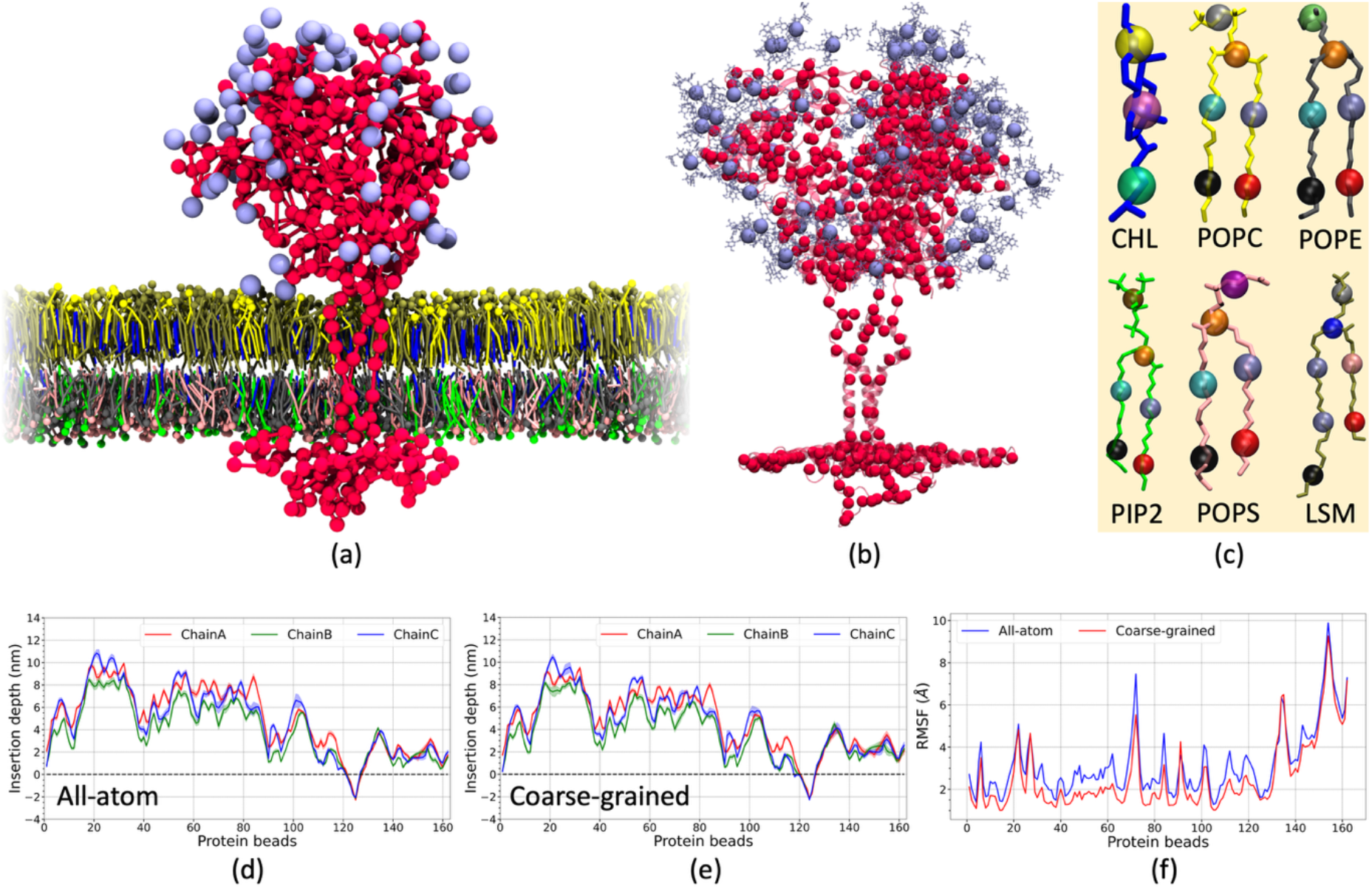
(a) Side view of the CG model of glycosylated Env embedded in an asymmetric bilayer containing POPC (yellow), LSM (tan), and cholesterol (blue) in the exofacial leaflet, and POPE (gray), POPS (pink), PIP2 (green), and cholesterol in the cytofacial leaflet. Env and glycans are represented in red and ice blue, respectively. (b) Pictorial representation showing the CG mapping of the glycosylated Env protein from the AA structure. (c) Schematic representation of the CG lipid model. All lipids were uniquely represented by 14 CG bead types assigned based on their chemical structures.^32^ Average depth of Env insertion into the membrane obtained from (d) AA and (e) CG simulations. (f) Root mean square fluctuation (RMSF) of Env obtained from AA and CG simulations.

The CG mapping operator for Env was constructed using the essential dynamics coarse-graining (ED-CG) method,^55^ which employs as input principal component analysis information obtained from AA MD simulations. Env was mapped onto 162 CG beads, with a resolution of approximately five amino acids per CG bead. Each glycan molecule was mapped onto a single CG bead (**Table S1**). The intra-protein interactions were modeled using a heteroelastic network model,^56^ in which the force field parameters were systematically optimized to match the fluctuation dynamics of the AA reference protein trajectory (see **Methods**). Solvent-free inter-protein interactions were modeled using screened electrostatics represented by a Yukawa potential. To model glycan-glycan interactions, we calculated the potential of mean force (PMF) governing glycan dimerization along the center-of-mass distance between glycans as the collective variable using AA simulations (**Figure S3**). The resulting PMF was used to derive the CG force field for glycan interactions (see **Methods**). An initial set of protein-lipid nonbonded parameters was obtained by analyzing pairwise radial distribution functions. These parameters were subsequently refined using the REM protocol (**Figure S4**). Previous studies have shown that, during REM iterations, CG simulations starting from random initial structures lead to improved results.^32, 57^ Therefore, to construct a robust protein-lipid model, we performed REM iterations starting from a random lipid distribution around the protein rather than from a CG-mapped AA simulation frame.

Following construction of the CG force field for the Env-lipid ensemble, we simulated glycosylated Env embedded in a HIV-1 mimetic asymmetric bilayer (**Figure 2a**) using the same membrane composition as in the AA simulations. From these CG simulations, we calculated the RMSF and insertion depth of Env in the membrane. The results are shown in **Figure 2**. The results show that the CG model accurately captures the protein dynamics in the membrane. The MPER was observed to interact at the interface of water and the exofacial leaflet, whereas the CT domain interacts at the cytofacial leaflet-water interface, as observed in the AA simulations. The CG model also captures the characteristic tilting motion of the Env ectodomain with respect to the membrane normal observed in AA simulations (**Figure S5**). Overall, the bottom-up CG model was found to accurately reproduce the Env-lipid interactions at the CG level observed in the reference AA simulations napped to the same resolution.

### Env proteins form clusters in the native viral envelope

HIV-1 expresses 7 to 14 Env proteins on its surface.^13, 14^ Despite having the same number of Env proteins, immature or partially mature virions show reduced infectivity compared to mature virions due to differences in their lateral organization.^20, 21^ In an immature virion, the Gag lattice underneath the membrane restricts the movement of Env proteins. Eggeling and coworkers showed that Env in immature virions exhibits significantly slower mobility compared to that in mature virions.^23^ However, a CT-truncated (ϕCT) variant of Env displays similar mobility in both mature and immature virions, which is also comparable to that of full-length Env on the mature virion surface.^23^ Kräusslich and coworkers have shown that Env proteins are randomly distributed in immature virions but predominantly form a cluster in mature virions.^20^ They also demonstrated that the CT domain plays an important role in Env clustering. The efficacy of viral entry was found to be directly correlated with the propensity for Env clustering. Maturation-induced Gag cleavage was shown to render Env more mobile, leading to Env oligomerization.^20^ Using cryo-ET, Lee and coworkers observed MA domain density underneath Env in immature virions, but not in mature virions.^58^ Importantly, no maturation-induced structural changes in the Env ectodomain were observed. They hypothesized that in immature virions, the MA domain of the Gag polyprotein interacts with Env, restricting its mobility. Upon maturation, the SP2 peptide binds to MA to form the mature MA lattice,^59^ and Env is released from its interaction with MA. Then, the free Env on the mature virion surface eventually forms clusters that are important for receptor-mediated viral entry.

To study Env dynamics in a mature virion, we first simulated an entire HIV-1 envelope with a diameter of 80 nm using the bottom-up CG model (**Figure 3a**). The envelope was modeled as an asymmetric bilayer with the same composition as the AA bilayer, consisting of 83,000 lipids. Fourteen Env proteins were placed randomly on the surface of the envelope. This simulation setup accurately represents Env organization in the mature viral envelope, as discussed by Lee and coworkers.^58^ The overall protein-lipid ensemble consisted of approximately 460,000 CG beads. An excluded volume interaction was placed inside the virion to mimic the densely packed interior environment of a real mature virion. An equivalent system at AA resolution would contain approximately 100 million atoms. The timescale required to study Env aggregation across the entire virion surface is likely impossible to achieve using present day AA simulations due to the prohibitive computational cost. Therefore, we performed two independent replicate CG simulations, each for 200 × 10^6^ CG timesteps. We note that CG simulations are effectively accelerated, and therefore the CG simulation time represents a much longer effective timescale compared to AA simulations. During the simulations, Env clustering was observed on the viral envelope. However, formation of a single Env cluster was not achieved, as observed in the experimental studies,^20^ likely due to finite simulation time.

**Figure 3.**
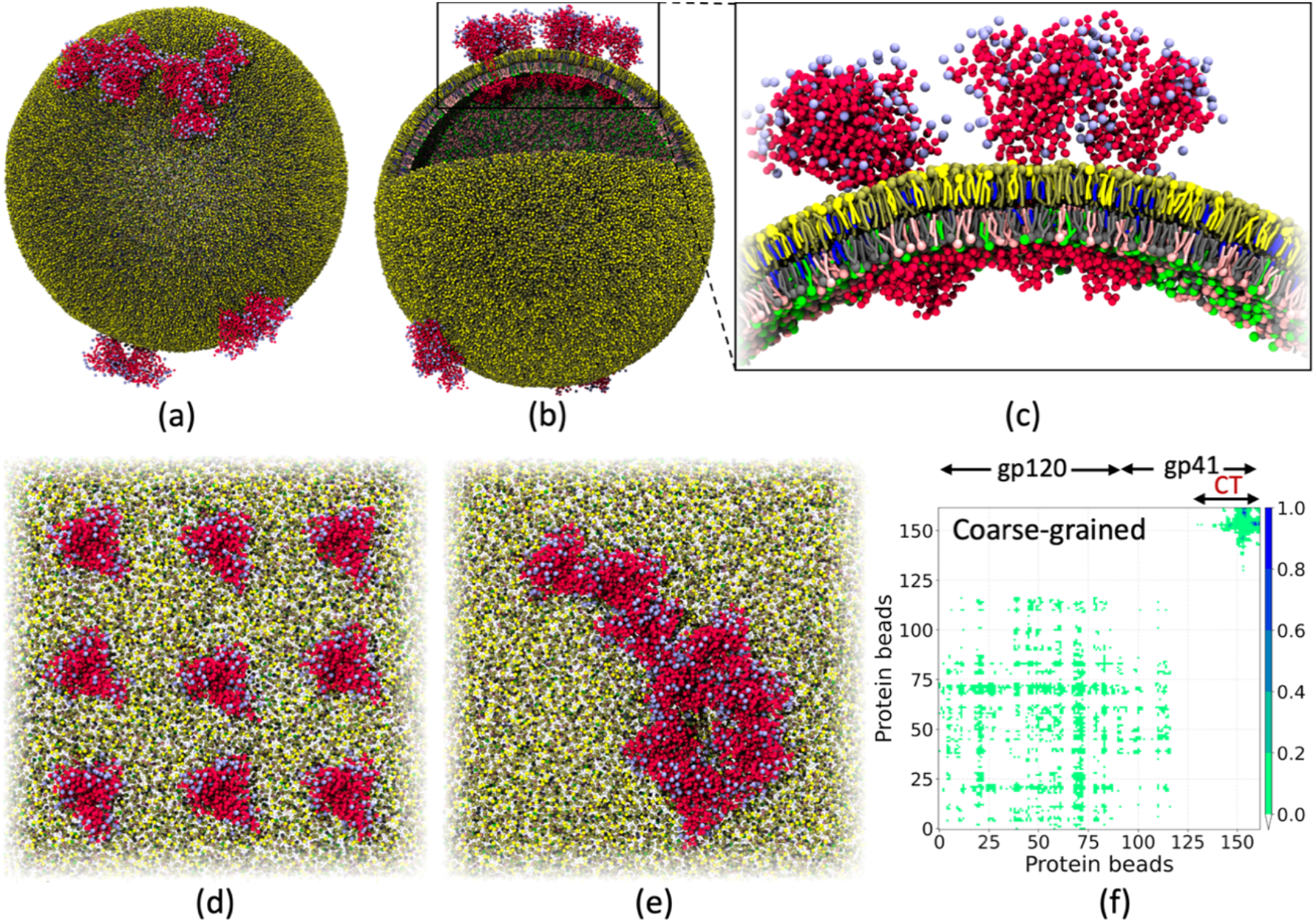
(a) Final configuration of the full virion envelope model containing 14 Env proteins obtained from the CG simulation. (b) Sliced view showing the interior (cytofacial leaflet), Env insertion, and membrane bilayer organization of the full envelope model. (c) Zoomed-in snapshot showing the asymmetric lipid organization in the full virion envelope. (d) Initial configuration and (e) final configuration from one of the replicate CG simulations, (f) contact map describing interactions between Envs obtained from CG simulations of 9 Envs in flat asymmetric bilayers. Env, glycans, POPC, LSM, and cholesterol, POPE, POPS, PIP2 are represented by red, ice blue, yellow, tan, blue, gray, pink, and green, respectively.

We also simulated a full HIV-1 envelope model starting from the same initial conformation, but with protein-protein interactions switched off while keeping all other CG force field parameters unchanged, to assess the importance of protein-protein interactions in Env clustering. During this simulation, we observed a scattered distribution of Env proteins (**Figure S6**). These results suggest that protein-protein interactions largely govern the Env clustering on the viral envelope.

To better sample the Env dynamics in a mature virion, we then simulated nine Env proteins in a HIV-1 mimetic flat asymmetric bilayer containing about 16,000 lipids using the CG force field. All Env proteins were initially placed separately from each other at the start of the simulations (**Figure 3d**). We performed three independent replica CG simulations, each for 200 × 10^6^ CG steps. During the simulations, we observed oligomerization of Env proteins, as well as the formation of a stable single Env cluster, consistent with previous experimental studies (**Figures 3e, S7, and S8**).^20^

### PIP2 accumulates near Env clusters through preferential interactions with the CT domain

We have previously studied^40^ the membrane-interacting domain of gp41 using AA simulations and showed that PIP2 lipids interact with the basic residues of the CT domain, leading to lipid demixing and PIP2 accumulation around the CT baseplate. Previous studies have shown that interactions between the CT domain and PIP2 lipids are crucial for the Env incorporation into the virion.^16, 60, 61^ Kräusslich and coworkers have shown that deletion of the CT domain or depletion of PIP2 lipids abrogates Gag’s influence on Env recruitment to the virion.^16^ They also hypothesized that membrane microdomains may play an important role in Env incorporation.

In our simulations, we observed accumulation of PIP2 lipids around the CTD in the AA simulations of full-length glycosylated Env (**Figure 1e**). We calculated the contact fraction describing interactions between Env proteins and PIP2 lipids from both AA and CG simulations (**Figure S9**). The results show that the CG model accurately reproduces the Env-PIP2 interaction patterns observed in the AA simulations, reflecting the value of a bottom-up CG’ing approach. We also calculated the PIP2 lipid density in the cytoplasmic leaflet in CG simulations of flat asymmetric bilayers containing nine Env proteins. The results are shown in **Figure 4**. We observe PIP2 sequestration around the Env clusters in all replicate simulations (**Figure S10**). These results suggest that PIP2 lipids likely play an important role in modulating the local membrane environment that facilitates Env recruitment and incorporation.

**Figure 4.**
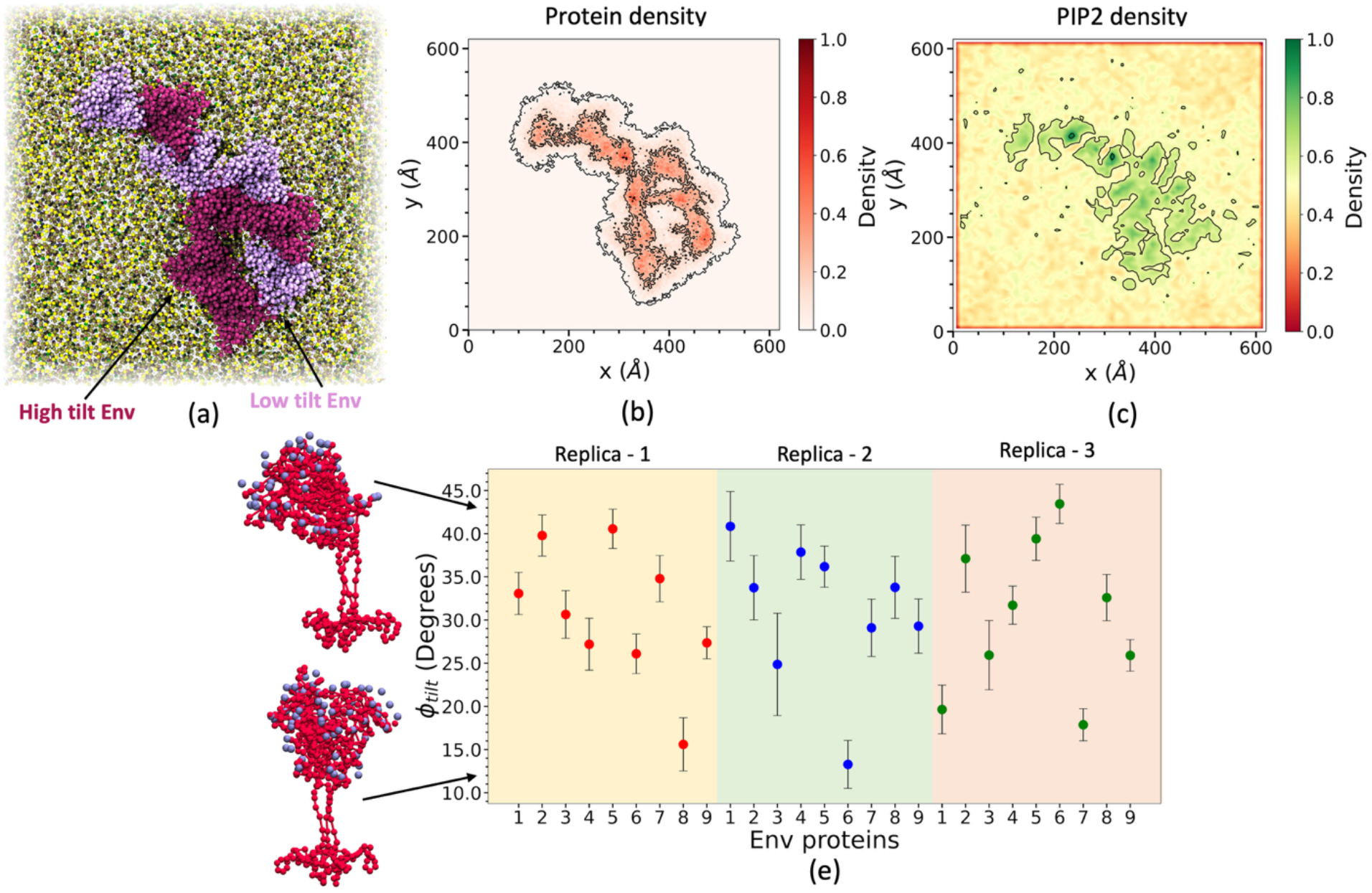
(a) Schematic representation showing high-tilt (purple) and low-tilt (ice-blue) Env conformations obtained from the CG simulations. (b) Env protein density and (c) PIP2 lipid density obtained from one of the replicate CG simulations of 9 Envs in flat asymmetric bilayers. (d) Tilt angle distribution of individual Envs obtained from three independent replicate simulations of Envs embedded in a flat asymmetric bilayer using the CG force field.

Croft et al.^45^ have characterized the tilt angle distributions of Env clusters in the viral membrane and showed that Env adopts both high-tilt and low-tilt conformations within a cluster.^45^ We characterized the tilt angle distribution of Env monomers and clusters in the viral membrane using CG simulations. The tilt angle distribution of Env proteins within clusters is shown in **Figure 4e**. While Env monomers in the CG simulations predominantly adopt a tilt angle of ∼30° (**Figure S5**), clustered Envs exhibit more diverse tilting orientations. Some Env ectodomains adopt highly tilted orientations, whereas others adopt more upright conformations (**Figures 4a and S11**), consistent with observations from cryo-ET experiments. This diversity in the tilting orientations of the Env ectodomain on the mature virion surface likely plays an important role in modulating the exposure of host receptor binding sites.

### Backmapping from CG to AA emphasizes that interactions between Env proteins are mediated by the CT domain

To characterize the interactions leading to Env clustering in CG simulations, we constructed a contact map by analyzing inter-protein CG bead distances (**Figure 3f)**. The contact map shows specific interactions between the CT domains, whereas interactions between the Env ectodomains appear scattered. To assess the importance of the CT domain in Env clustering, we further performed AA simulations of an Env dimer embedded in a HIV-1 mimetic asymmetric bilayer.

The artificial intelligence (AI) machine learning-based MSBack protocol was used to backmap the Env dimer structure from the CG simulation to AA resolution.^62^ The Env structure obtained from the AA simulations served as the reference state for the MSBack backmapping pipeline. The individual CG coordinates of the target Env dimer complex were used to fit the reference AA Env structure. The Env-Env interface was refined by perturbing the reference AA structure with Chroma to generate a backbone matching the target CG coordinates.^63^ Side chains were added using FlowBack,^64^ and the protein structure was relaxed using the mstool protocol.^65^ The resulting AA Env dimer structure was then embedded in a pre-equilibrated asymmetric bilayer and solvated with water and ions to generate the starting configuration for the simulations (**Figure 5a**). The AA system consisted of ∼3 million atoms. Two independent replicate production runs were performed, yielding a total of 1.2 μs of simulation time. The contact map defining interactions between Env protomers obtained from the AA simulations is shown in **Figure S12**. The results indicate that Env interactions are predominantly mediated by the CT domain, consistent with the CG simulations. Interactions between the CT domains observed in the Env dimer AA simulations are shown in **Figure 5b**. The backmapped AA simulations predict that basic residues in the C-terminal region of the CT domain play an important role in stabilizing the Env dimer structure. A previous study showed that mutation of Arg to Gln can modulate electrostatic interactions in membrane proteins without significantly affecting their structure or thermal stability.^66^ Based on this, we propose that R845Q, R846Q, and R853Q mutants of Env could be studied to further assess the role of the CTD in Env oligomerization. Overall, the results presented in this section highlight the power of the combined CG to backmaped AA simulation approach, enabled by AI technology.

**Figure 5.**
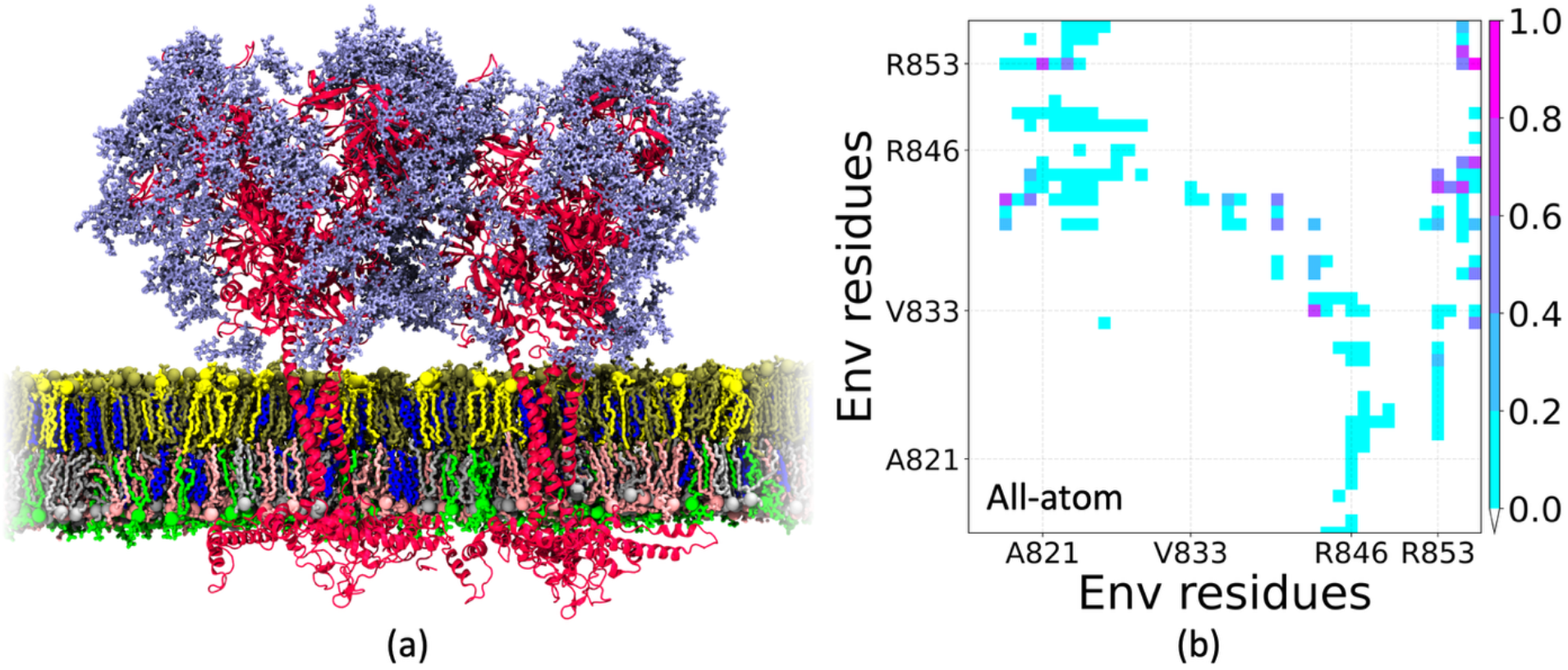
(a) Side view of the AA structure of glycosylated Env dimer obtained from backmapped CG simulations embedded in an asymmetric membrane, and (b) contact map describing interactions between Envs (CT domain) obtained from AA simulations of Env dimer. Env, glycans, POPC, LSM, and cholesterol, POPE, POPS, PIP2 are represented by red, ice blue, yellow, tan, blue, gray, pink, and green, respectively.

### Env ectodomain epitopes remain available in mature virions

Broadly neutralizing antibodies (bNAbs) that efficiently neutralize HIV-1 target Env trimers on the surface of the virion. bNAbs can be categorized into five different classes based on the positions of their target epitopes on Env: the CD4 binding site, the V1/V2 loop and N160 glycan, the V3 region and N332 glycan, the gp120-gp41 interface, and the MPER-directed antibodies.^67-69^ Chen and coworkers showed that antibodies targeting the CD4 binding site, V1/V2 loop, and V3 loop can effectively bind to the prefusion conformation of Env, whereas MPER-directed antibodies do not bind to the prefusion conformation and instead target fusion-intermediate configurations of Env.^70, 71^ Im and coworkers studied epitope exposure on the prefusion configurations of glycosylated Env monomer using AA MD simulations and showed that the MPER region is largely occluded for antibody binding, whereas antibodies can bind efficiently to the V3 loop.^42^

We analyzed the accessibility of antibodies from different classes to their epitopes on Env. Specifically, we studied antibody 35022 targeting the gp120-gp41 interface, VRC01 targeting the CD4 binding site, 10E8 targeting the MPER, PG9 targeting the V1/V2 loops, and PGT128 targeting the V3 loop. To assess the accessibility of the antibody binding sites, we selected experimental structures of the antibodies bound to their corresponding epitopes. The epitope structures were then RMSD fitted to the Env CG structures obtained from the simulations. An epitope was considered occluded if more than three CG bead clashes were detected. We calculated the accessibility of each epitope by analyzing the conformation of each protomer in the trimeric Env structures. The results obtained from the CG simulations are shown in **Figure 6**. An accessibility score of unity signifies that the antibody can bind to the epitopes on all three protomers of Env in every frame of the CG simulations. Due to the clustering of Env proteins in mature virions, epitopes become more occluded compared to those in monomeric Env.

**Figure 6.**
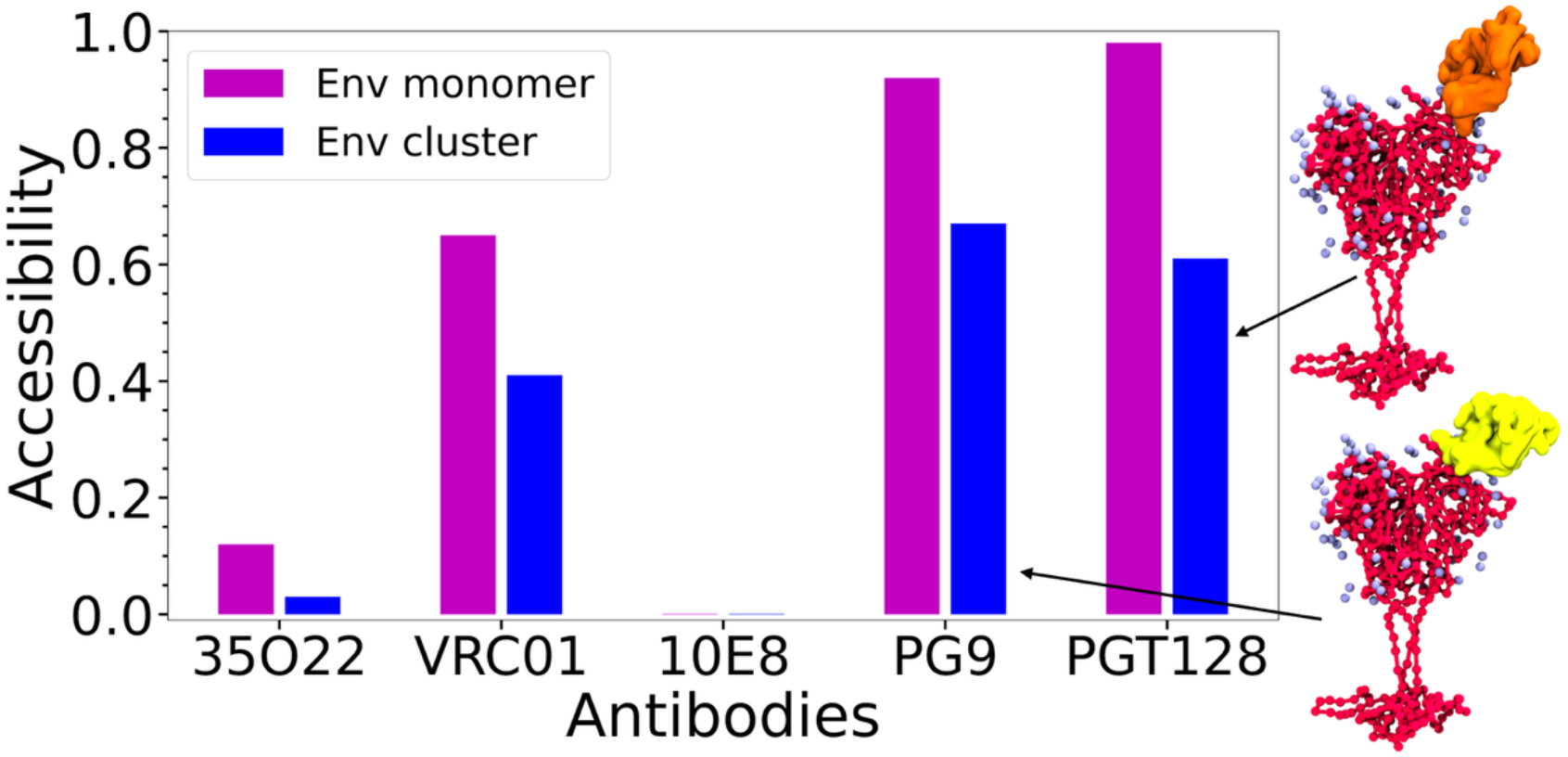
Accessibility of different antibodies to Env monomers and clusters. An accessibility value of 1 indicates that the epitope for that antibody remains available to each protomer in the trimeric Env structure throughout the simulation.

Although the 35022 antibody binds to the ectodomain of Env, tilting of Env causes the antibody to clash with the membrane, leading to a lower accessibility score. However, epitopes targeted by PG9 and PGT128, corresponding to the V1/V2 loops and the V3 loop, remain more accessible than the other epitopes in mature virions. The epitope of 10E8 (MPER) remains completely occluded in both Env monomers and clusters due to clashes with both the membrane and other protein residues, consistent with previous studies. From our simulations, we conjecture that antiviral designs based on the structures of PG9 or PGT128, specifically targeting the V1, V2, or V3 loops, may be more effective in targeting mature virions before they interact with receptors on healthy human cells.

Overall, the CG model of the full viral envelope provides a framework for large-scale simulations of HIV-1 particles at different stages of the viral life cycle, enabling a better understanding of virion organization and aiding in the development of strategies to prevent infection.

## Conclusions

The present study utilizes both AA and CG simulations synergistically to investigate the structural properties and dynamics of one or more HIV-1 Env proteins in viral membranes. Using AA simulations, we first simulated a fully glycosylated Env protein in HIV-1 mimetic asymmetric bilayers. The Env protein was observed to anchor to the bilayer through the C-terminal domain of gp41, where the MPER domain interacted with the exofacial leaflet-water interface. At the same time, the CT domain formed a stable baseplate around the TMD on the cytofacial side of the bilayer. The ectodomain of the Env protein was found to adopt a tilted conformation with respect to the membrane normal both in the presence and absence of its glycan shield. However, in the glycosylated Env, the N88, N611, and N625 glycans interact with lipids in the exofacial leaflet as a result of the tilting of Env.

We next constructed a bottom-up CG model of Env embedded in HIV-1 membranes using AA simulations as a reference. Assessment of the RMSF of the protein, protein insertion depth in the membrane, and protein-lipid contacts showed that the CG model accurately reproduces the properties of the AA model as projected onto a CG mapping. The CG model also captures the characteristic tilting motion of the Env ectodomain. We then performed CG simulations of multiple copies of Env by mimicking the Env environment in a mature HIV-1 virion envelope. The Env proteins were found to oligomerize, consistent with previous experimental studies.^20^ The PIP2 lipid sequestration was observed underneath Env clusters, where basic residues in the CTD interact with PIP2, leading to lipid demixing and accumulation of PIP2 around the CTD. Within clusters, Env proteins adopt both high-tilt and low-tilt conformations, consistent with previous experimental observations.^45^

Using AI technology, we then backmapped the CG Env dimer to AA resolution using the machine learning-based MSBack protocol and performed further AA simulations with the dimer embedded in an asymmetric bilayer. These simulations show that the CT domain plays an important role in Env clustering through interactions mediated by its C-terminal region. Additionally, we characterized the exposure of broadly neutralizing antibody epitopes on Env clusters on the viral envelope. Although maturation-induced Env clustering leads to more occluded epitope conformations, the V1/V2 and V3 loop regions of Env were found to remain accessible to antibodies in both monomers and clusters. Based on this observation, explicit modeling of bNAbs on Env, combined with efficient sampling of epitope and antibody conformations, could provide a better assessment of epitope accessibility.

Overall, this study characterizes maturation-induced Env dynamics on the viral envelope and provides insight into the interactions and conformational states that drive Env clustering. Understanding epitope exposure may help guide the design of more potent antivirals targeting mature HIV-1 particles. In future work, explicit models of the Gag polyprotein and capsid will be integrated into the full HIV-1 model to better represent additional aspects of the HIV-1 lifecycle.

## Methods

### Fully glycosylated Env construct

We constructed the full-length Env (gp120/gp41)_3_structure by combining the subtomogram structure of the ectodomain and the NMR structure of the MPER-TMD-CT domain (**Figure 1**). The trimeric ectodomain structure of Env was obtained from PDB ID 7SKA, which was resolved by analyzing Env bound in its native viral membrane,^33^ and the MPER-TMD-CT structure was obtained from PDB ID 7LOI, which is the only available full-length structure of the entire membrane-interacting domains of Env.^7^ In structure 7SKA, residues 653-662 located at the C-terminus are reported to adopt a loop structure. The trimeric ectodomain was attached at Leu660 to the N-terminus of structure 7LOI to construct the full-length trimeric Env structure. The missing residues in the protein construct were built using MODELLER.^72^ We modeled N-linked glycans at 24 glycosylation sites in each protomer (**Figure 1**). Previous experimental studies using mass spectrometry have characterized the glycan structures at different sites on Env.^34-37^ We selected the most probable glycan composition for each site that also matched the glycan fragments resolved in the ectodomain subtomogram structure. All glycans on Env were modeled using the CHARMM-GUI glycan builder (**Tables S1 and S2**).^73^

### All-atom simulation of Env in lipid bilayer

We simulated Env embedded in an asymmetric bilayer using the all-atom CHARMM36m force field.^43^ Env was simulated both with and without glycans. The composition of the asymmetric bilayer was chosen to mimic the HIV-1 virion membrane, where the exofacial leaflet is composed of 20 mol% POPC, 40 mol% LSM, and 40 mol% cholesterol, while the cytofacial leaflet is composed of 40 mol% POPE, 30 mol% POPS, 15 mol% PIP2, and 15 mol% cholesterol. The protein-embedded lipid bilayer was solvated using 175 waters per lipid, defined by the TIP3P water model,^74^ and 0.15 M KCl. The initial structures of the solvated protein-lipid systems were prepared using the CHARMM-GUI membrane builder.^75^ Each bilayer was equilibrated for 50 ns following the CHARMM-GUI protocol.^76^ Following equilibration, a 1-2 μs AA MD production run was performed (**Table S3**). The production run was carried out in the constant NPT ensemble using a leap-frog integration method with a time step of 2 fs. The simulation temperature was maintained at 303 K using the Nose-Hoover thermostat. Pressure during equilibration and production runs was maintained at 1 bar using semi-isotropic Berendsen and Parrinello-Rahman barostats, respectively. All simulations using the AA force field were performed using GROMACS.^77^

### Dimerization free energy of glycans using all-atom MD simulations

Umbrella sampling (US) simulations were performed to calculate the interaction energies of FA3, M8, and M5 glycans. First, two glycan molecules were placed in a cubic box of length 9 nm and solvated with water and 0.15 M KCl. The system was then equilibrated for 5 ns, followed by a 100 ns production run. To perform US, harmonic restraints of 500 kJ mol^-1^ nm^-2^ were applied at intervals of 0.15 nm along the center-of-mass (COM) distance between the glycans. Each umbrella window was simulated for 100 ns. The weighted histogram analysis method (WHAM) was used to unbias the simulations and obtain the final potential of mean force (PMF).^78^ The US simulations using the AA force field were performed using PLUMED^79^ patched with GROMACS.

### Construction of protein-lipid coarse-grained model

The CG representations of proteins and lipids were defined by coordinates *R*^*N*^ derived from the corresponding AA coordinates *r*^*n*^ using a mapping operator *M(r*^*n*^*) = R*^*N*^, where *N* and *n* are the numbers of CG and AA sites, respectively.^51, 52^

### Coarse-grained force field for lipids

Six CG beads were assigned to POPC, POPE, POPS, and PIP2, whereas cholesterol and LSM were mapped onto three and seven CG beads, respectively. Bond and angular force field of the lipids were obtained using the multiscale coarse-grained (MS-CG) force-matching approach.^29, 50-52^ The initial “solvent-free” nonbonded force fields of the lipids were also derived from the MS-CG approach and were further optimized using the relative entropy minimization (REM) protocol^54^ by minimizing relative entropy *S*_*rel*_ between CG and the corresponding AA reference model:

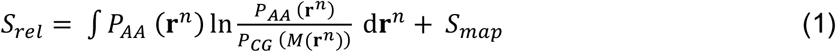

where *P*_*AA*_*(r*^*n*^) and *P*_*CG*_*(M(r*^*n*^)) represent the probability distribution of the AA and CG configurations, respectively, and *S*_*map*_is the average entropy that arises from the degeneracy of the mapping operator. In practice, the model parameters Ɵ were iteratively refined by minimizing *S*_*rel*_using a step function *λ* as

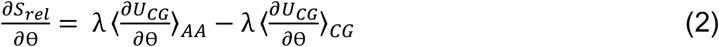

where *U*_*CG*_ is the CG force field. A detailed description of the method used to derive the lipid force field was provided in a previous study.^32^ CG model optimization was performed using the OpenMSCG software.^53^

### Coarse-grained force field for protein

We used the essential dynamics coarse-graining (ED-CG) approach to derive a CG mapping operator for Env.^55^ EDCG protocol uses AA protein trajectories as a reference and groups protein residues into CG beads such that the CG model captures the collective AA motions. The Env protein was projected onto 162 CG beads, corresponding to a resolution of ∼ 5 amino acids per CG bead, and each glycan was mapped onto a single CG bead. The overall CG protein force field of Env was defined as

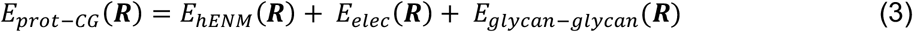

Intra-protein interactions were modeled using the hetero-elastic network model (hENM) with a cutoff of 20 Å, in which protein CG beads were connected by a bonded potential *K*_*ij*_ *(r*_*ij*_ *– r*_*0, ij*_)^*2*^, where *r*_*0,ij*_ is the equilibrium distance between CG bead *i* and *j*.^56^ Following the hENM protocol, *K*_*ij*_were iteratively optimized to match the protein dynamics observed in the reference AA simulation.

Inter-protein interactions consisted of screened electrostatic potential (*E*_*elec*_) and glycan-glycan interaction potential (*E*_*glycan-glycan*_). For *E*_*elec*_, a Yukawa potential was used, with the inverse Debye length set to 0.1274 Å^-1^, and an effective dielectric constant was set to 8. Glycan-glycan interactions were parameterized by fitting the PMF obtained from AA simulations to a pairwise Gaussian potential 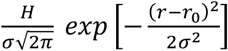. We calculated the PMFs defining binding interaction for FA3, M8, and M5 glycans. Glycans FA3G3, FA3, and FA2, and glycans M9, M8, and M7 were modeled using the same parameters. Inter-glycan interactions were defined as 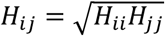 and *r*_*0,ij*_=(*r*_*0,ij*_+*r*_*0,ij*_)^2^

### Coarse-grained force field for protein-lipid

Protein-lipid nonbonded interactions were modelled using pairwise Gaussian potentials and excluded volume interactions. The excluded volume parameters were determined by analyzing protein-lipid radial distribution function. Initial parameters for the nonbonded interactions were derived using the Boltzmann-Inverse free energy profiles. The parameter *H*_*ij*_ for each protein-lipid pair was then iteratively optimized using the REM protocol (**eq 2**) to obtain the final protein-lipid force field.

### Coarse-grained simulations of Env in lipid bilayers

Initial configurations of the CG model of Env embedded lipid bilayers were prepared using PACKMOL.^80^ All protein-lipid systems were simulated using a time step of 15 fs. For constant NVT simulations, a Langevin thermostat with a coupling constant of 10 ps was used to maintain the temperature at 303 K. For constant NPT simulations, an additional barostat was applied along the *xy*-dimension (parallel to the membrane surface) to maintain a pressure of 0 bar. A 25 Å cutoff distance was used for all nonbonded interactions between CG beads. The protein-lipid systems simulated using the CG force field are listed in **Table S3**. All CG simulations were performed using LAMMPS.^81^

See **Data Availability** for all parameters in the model.

## Supporting Information

The supplementary information contains (**Table S1 and S2**) Glycan structures and associated CHARMM scripts; (**Table S3**) Systems simulated in this study; (**Figure S1**) Contact map of glycans and lipids; (**Figure S2**) Probability density of hinge angles; (**Figure S3**) PMFs govorning glycan dimerization; (**Figure S4**) REM iterations for protein-lipid force field; (**Figure S5**) Env tilt angle in CG simulations; (**Figure S6**) CG simulation of full virion; (**Figure S7**) Env clustering in CG simulations; (**Figure S8**) CG simulations of flat bilayer; (**Figure S9**) Interactions between CT domain and PIP2/POPS; (**Figure S10**) Env and PIP2 density in CG simulations; (**Figure S11**) Env tilting in CG simulations; (**Figure S12**) Contact map of Env dimer in AA simulations.

## Data Availability

The initial and final structures from three AA replica simulations, along with AA model parameters of HIV-1 Env monomers and dimers embedded in asymmetric membrane bilayers composed of POPC, LSM, cholesterol, POPS, POPE, and PIP2, are provided. GROMACS input files used to perform AA equilibration and production runs are also included. In addition, CG model parameters for Env proteins embedded in an HIV-1 mimetic bilayer, LAMMPS input files used to perform CG simulations of flat bilayers and full virion envelopes containing multiple Env proteins, and all CG trajectories are available at: https://doi.org/10.5281/zenodo.20261483.

## Acknowledgments

This research was supported by the National Institute of Allergy and Infectious Diseases (NIAID) of the National Institutes of Health (NIH) through grant R01AI178850. The content is solely the responsibility of the authors and does not necessarily represent the official views of the National Institutes of Health. The authors gratefully acknowledge computational resources provided by the University of Chicago Research Computing Center and the Frontera supercomputer at the Texas Advanced Computer Center funded by the National Science Foundation (OAC-1818253).

## TOC Graphic

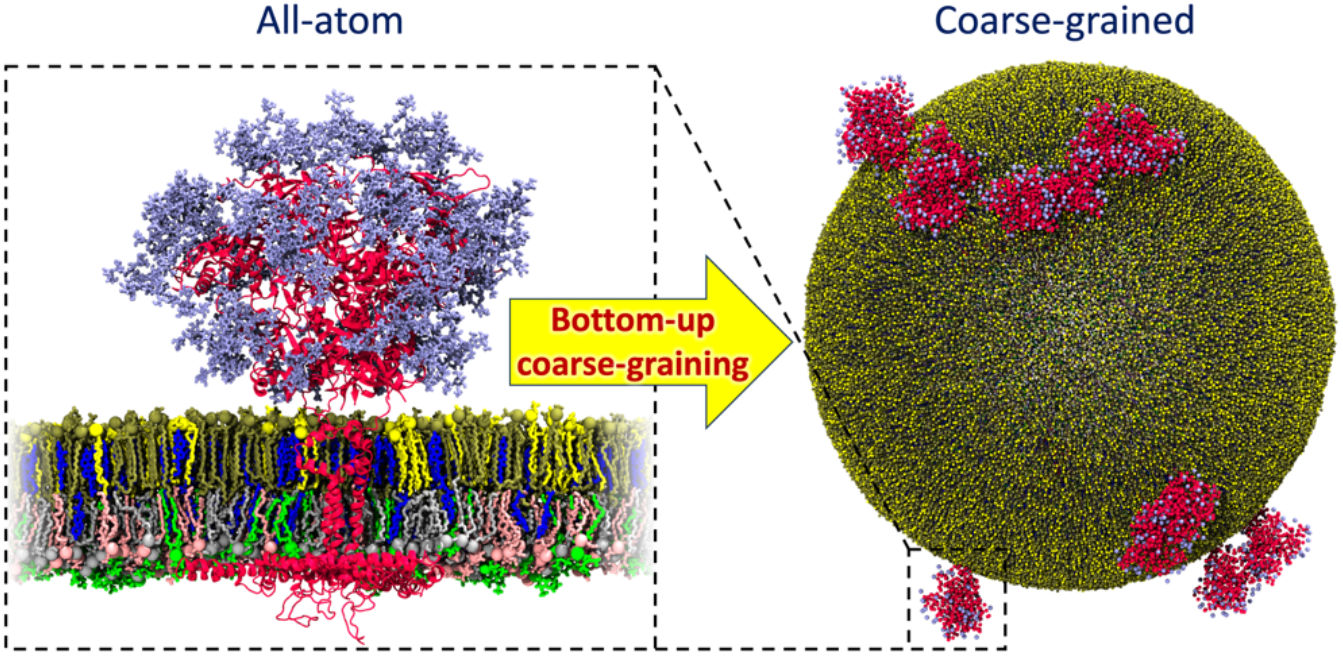

