## Supplementary Information for "Structure and Dynamics of the HIV-1 Envelope Protein on the Virion Envelope"

**Table S1:** Glycan structures simulated in this study.

| Position | Glycan | Structure |
|---|---|---|
| 88 | FA2 | 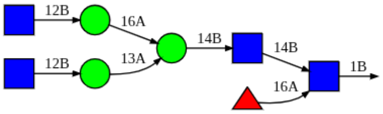 |
| 136 | FA2 | 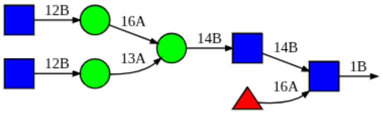 |
| 156 | M9 | 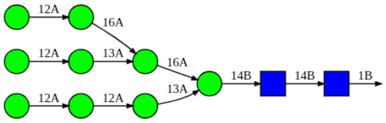 |
| 160 | M8 | 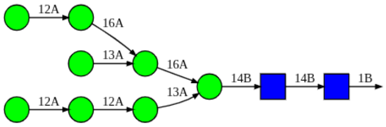 |
| 188 | FA2 | 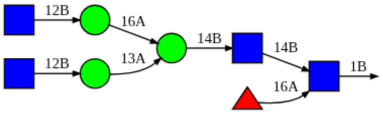 |
| 197 | M8 | 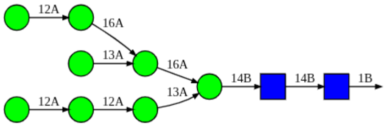 |

| | | |
|---|---|---|
| 234 | M9 | 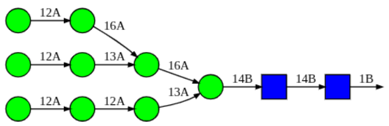 |
| 262 | M9 | 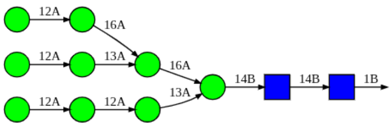 |
| 276 | M7 | 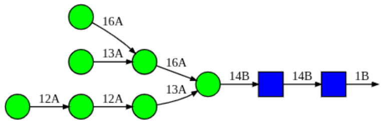 |
| 295 | M9 | 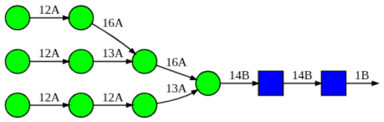 |
| 301 | M9 | 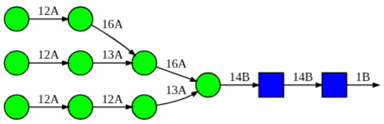 |
| 331 | M9 | 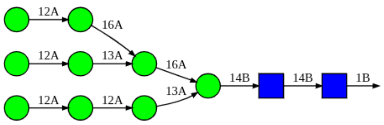 |
| 338 | M9 | 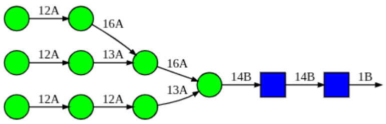 |
| 355 | M5 | 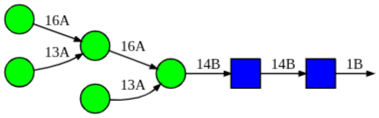 |
| 361 | M9 | 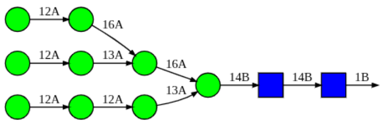 |
| 385 | M9 | 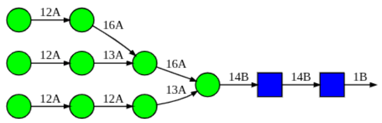 |

| | | |
|---|---|---|
| 391 | M9 | 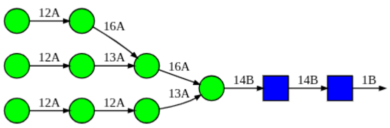 |
| 397 | M8 | 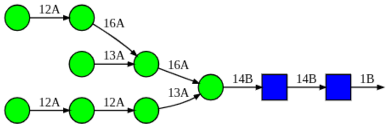 |
| 406 | FA2 | 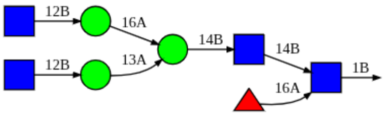 |
| 411 | M8 | 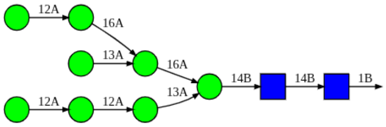 |
| 461 | FA3 | 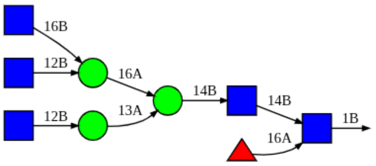 |
| 611 | FA3 | 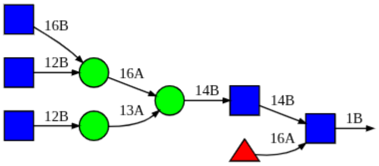 |
| 625 | FA2 | 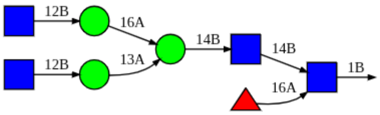 |
| 637 | FA3G3 | 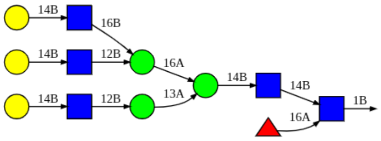 |

**Table S2:** CHARMM script used to generate glycan structures simulated in this study.

| Glycan | Structure | CHARMM script |
|---|---|---|
| FA3G3 | 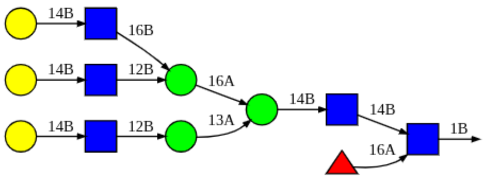 | 1 BGLCNA<br>2 - 14B: BGLCNA<br>3 - - 14B: BMAN<br>4 - - - 16A: AMAN<br>5 - - - - 16B: BGLCNA<br>6 - - - - - 14B: BGAL<br>7 - - - - 12B: BGLCNA<br>8 - - - - - 14B: BGAL<br>9 - - - 13A: AMAN<br>10 - - - - 12B: BGLCNA<br>11 - - - - - 14B: BGAL<br>12 - 16A: AFUC |
| FA3 | 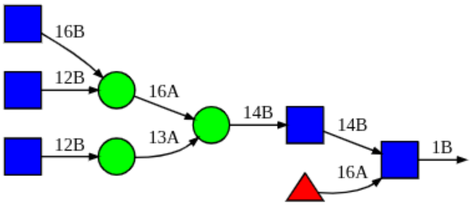 | 1 BGLCNA<br>2 - 14B: BGLCNA<br>3 - - 14B: BMAN<br>4 - - - 16A: AMAN<br>5 - - - - 16B: BGLCNA<br>6 - - - - 12B: BGLCNA<br>7 - - - 13A: AMAN<br>8 - - - - 12B: BGLCNA<br>9 - 16A: AFUC |
| FA2 | 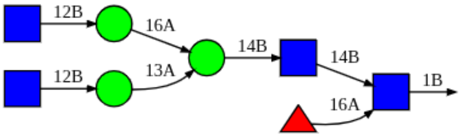 | 1 BGLCNA<br>2 - 14B: BGLCNA<br>3 - - 14B: BMAN<br>4 - - - 16A: AMAN<br>5 - - - - 12B: BGLCNA<br>6 - - - 13A: AMAN<br>7 - - - - 12B: BGLCNA<br>8 - 16A: AFUC |
| M9 | 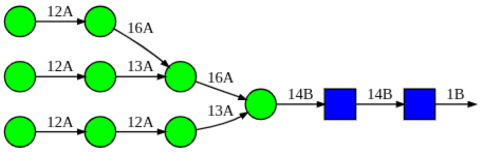 | 1 BGLCNA<br>2 - 14B: BGLCNA<br>3 - - 14B: BMAN<br>4 - - - 16A: AMAN<br>5 - - - - 16A: AMAN<br>6 - - - - - 12A: AMAN<br>7 - - - - 13A: AMAN<br>8 - - - - - 12A: AMAN<br>9 - - - 13A: AMAN<br>10 - - - - 12A: AMAN<br>11 - - - - - 12A: AMAN |

| | | |
|---|---|---|
| M8 | 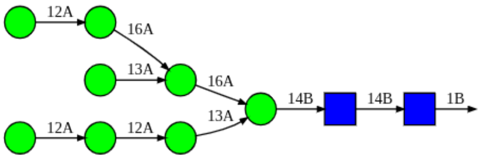 | 1 BGLCNA<br>2 - 14B: BGLCNA<br>3 - - 14B: BMAN<br>4 - - - 16A: AMAN<br>5 - - - - 16A: AMAN<br>6 - - - - - 12A: AMAN<br>7 - - - - 13A: AMAN<br>8 - - - 13A: AMAN<br>9 - - - - 12A: AMAN<br>10 - - - - - 12A: AMAN |
| M7 | 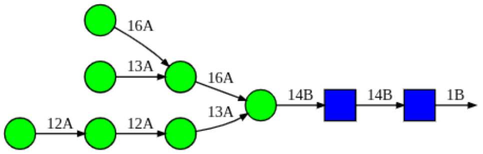 | 1 BGLCNA<br>2 - 14B: BGLCNA<br>3 - - 14B: BMAN<br>4 - - - 16A: AMAN<br>5 - - - - 16A: AMAN<br>6 - - - - 13A: AMAN<br>7 - - - 13A: AMAN<br>8 - - - - 12A: AMAN<br>9 - - - - - 12A: AMAN |
| M5 | 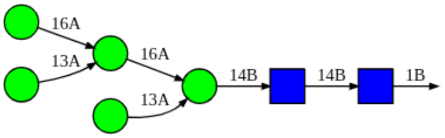 | 1 BGLCNA<br>2 - 14B: BGLCNA<br>3 - - 14B: BMAN<br>4 - - - 16A: AMAN<br>5 - - - - 16A: AMAN<br>6 - - - - 13A: AMAN<br>7 - - - 13A: AMAN |

**Table S3**: Total simulation times performed for each membrane system studied using AA and CG force fields. We note that CG simulations are effectively accelerated, and therefore the CG simulation time represents a much longer effective timescale compared to AA simulations.

| System | Replica | Total simulations |
|---|---|---|
| **All-atom** | | |
| Env with glycans in an asymmetric bilayer | 3 | 4 μs |
| Env without glycans in an asymmetric bilayer | 3 | 3 μs |
| Env dimer with glycans in an asymmetric bilayer | 2 | 1.2 μs |
| **Coarse-grained** | | |
| One Env in flat asymmetric bilayer | 2 | $200 \times 10^6$ CG steps |
| 9 Env in flat asymmetric bilayer | 3 | $600 \times 10^6$ CG steps |
| 14 Env in full virion envelope model | 2 | $400 \times 10^6$ CG steps |
| 14 Env in full virion envelope model (no protein-protein interactions) | 1 | $200 \times 10^6$ CG steps |

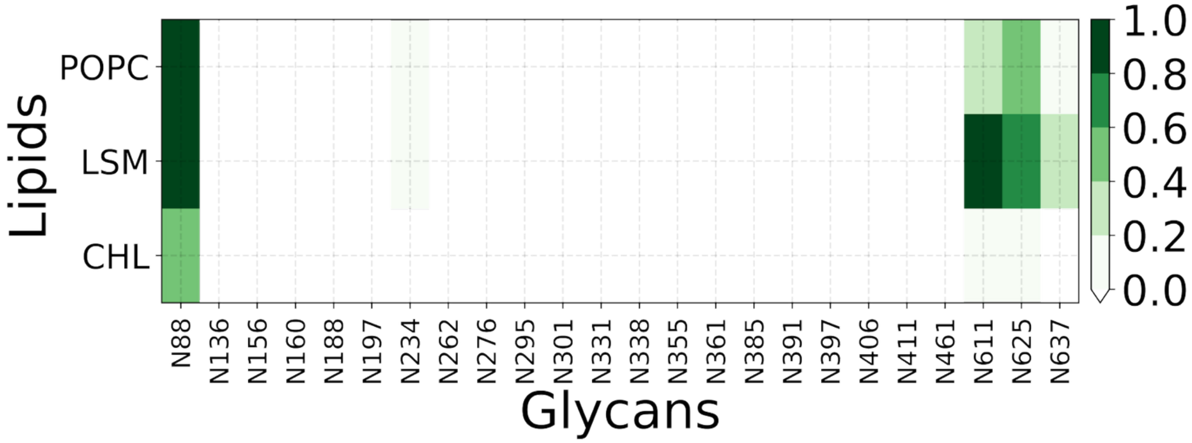


**Figure S1:** Contact map describing interactions between glycans and lipids obtained from AA simulations of glycosylated Env in asymmetric bilayers.

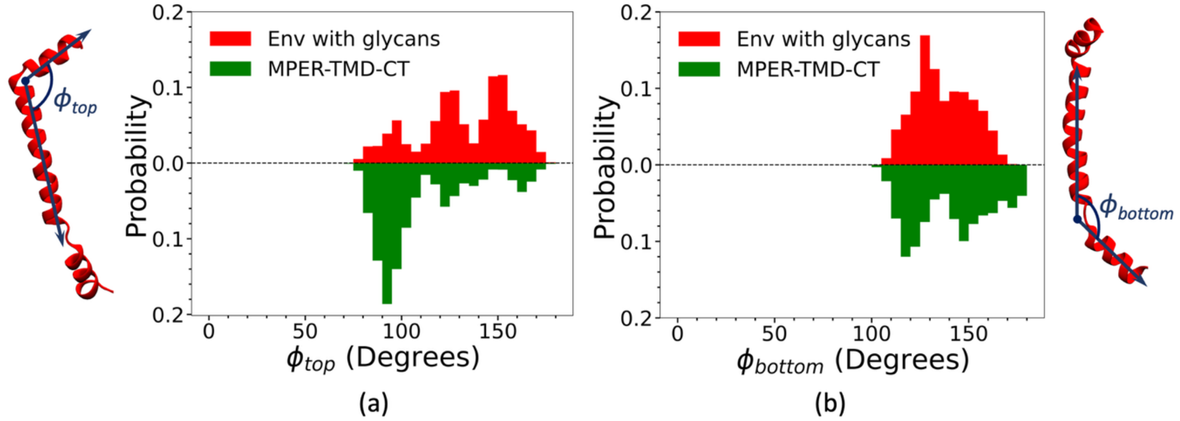


**Figure S2:** Probability density of the top ($\phi_{top}$) and bottom ($\phi_{bottom}$) hinge angles characterizing the structural ensemble of MPER-TMD obtained from AA simulations of glycosylated Env and MPER-TMD-CT trimers.

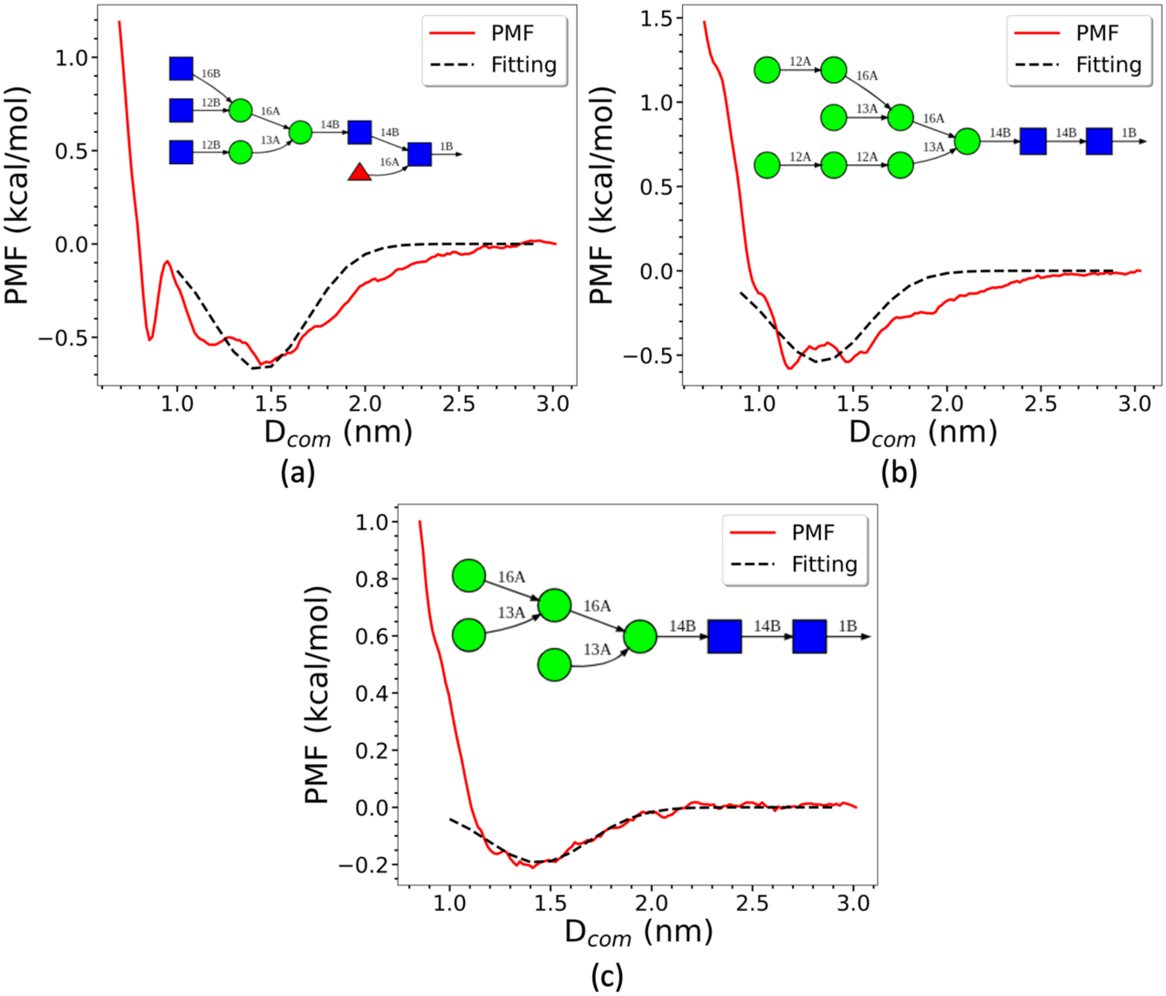


**Figure S3:** Potential of mean force (PMF) governing dimerization of (a) FA3, (b) M8, and (c) M5 glycans calculated along the center-of-mass (COM) distances between the glycans, obtained from AA simulations. The PMF and the Gaussian fit are shown in red and black, respectively.

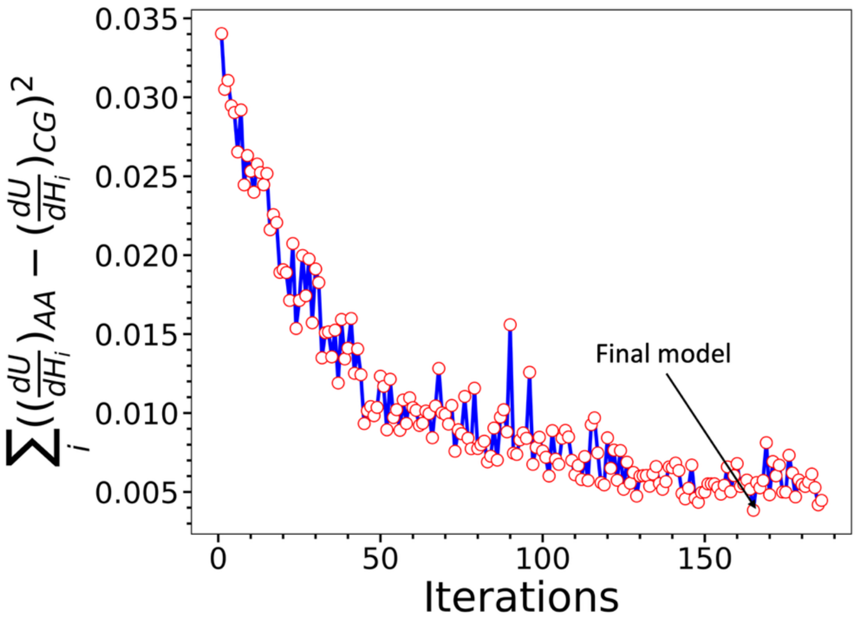


**Figure S4:** Squared differences in the gradients of force field with respect to the model parameters obtained by analyzing AA and CG simulations during the REM iterations used to refine CG protein-lipid force field.

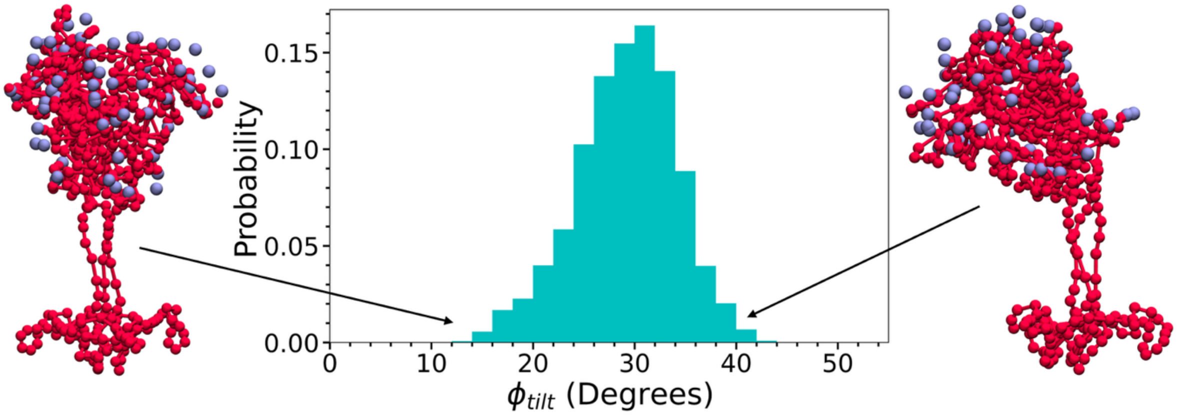


**Figure S5:** Tilt angle distribution of the Env ectodomain relative to the membrane normal obtained from the CG simulation.

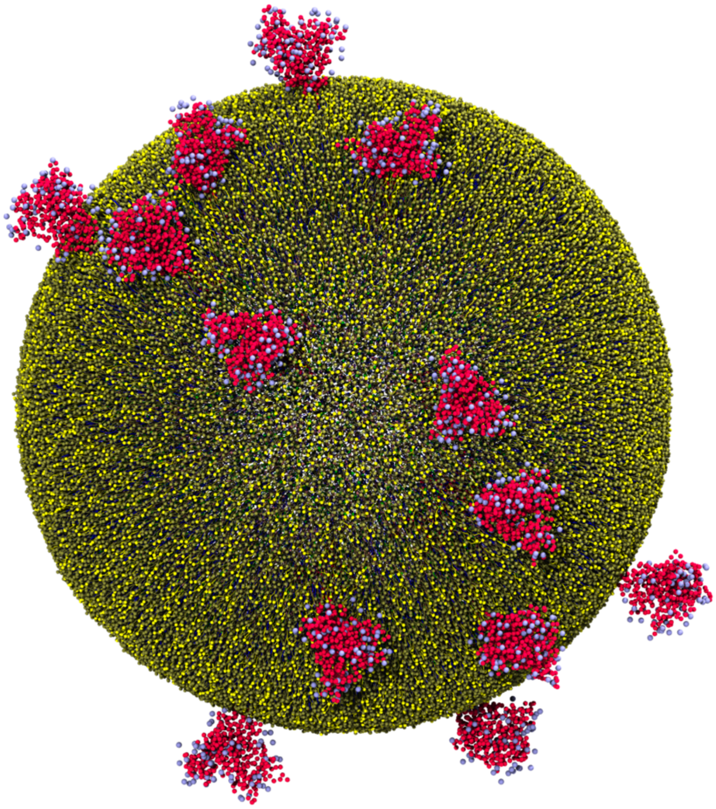

**Figure S6:** Final configuration of the full virion envelope model containing 14 Env proteins obtained from the CG simulation. In this case, during the simulation the protein-protein interactions **were switched off** while keeping all other forcefield parameters the same. Note that the Env clustering is greatly reduced over the full model (figs 3a and 3b of the main text), thus emphasizing the role of the protein-protein interactions in their clustering behavior.

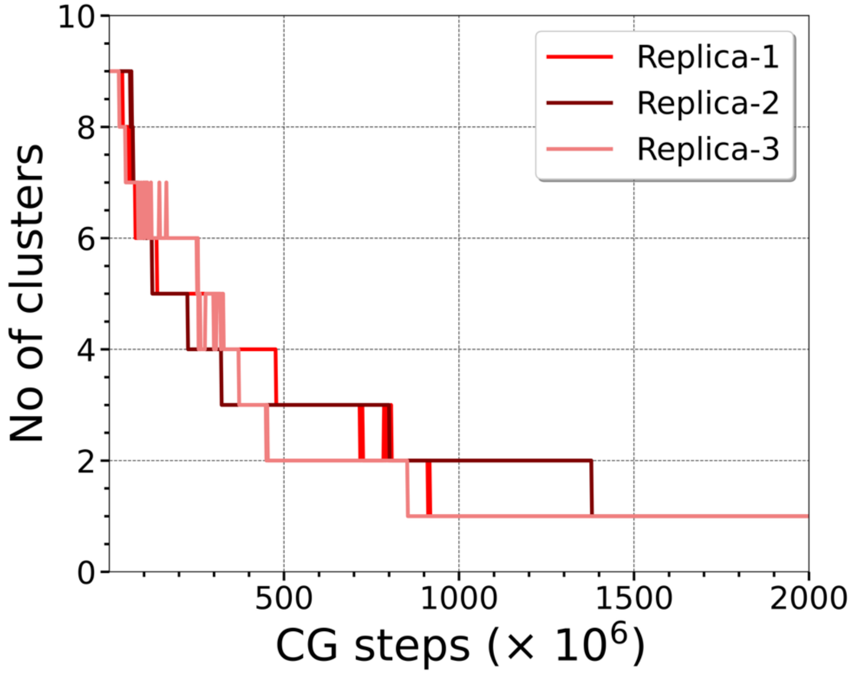


**Figure S7:** Number of Env clusters formed in CG simulations of nine Env trimers in flat asymmetric bilayers. Decrease in the number of Env clusters over time reflects progressive Env association.

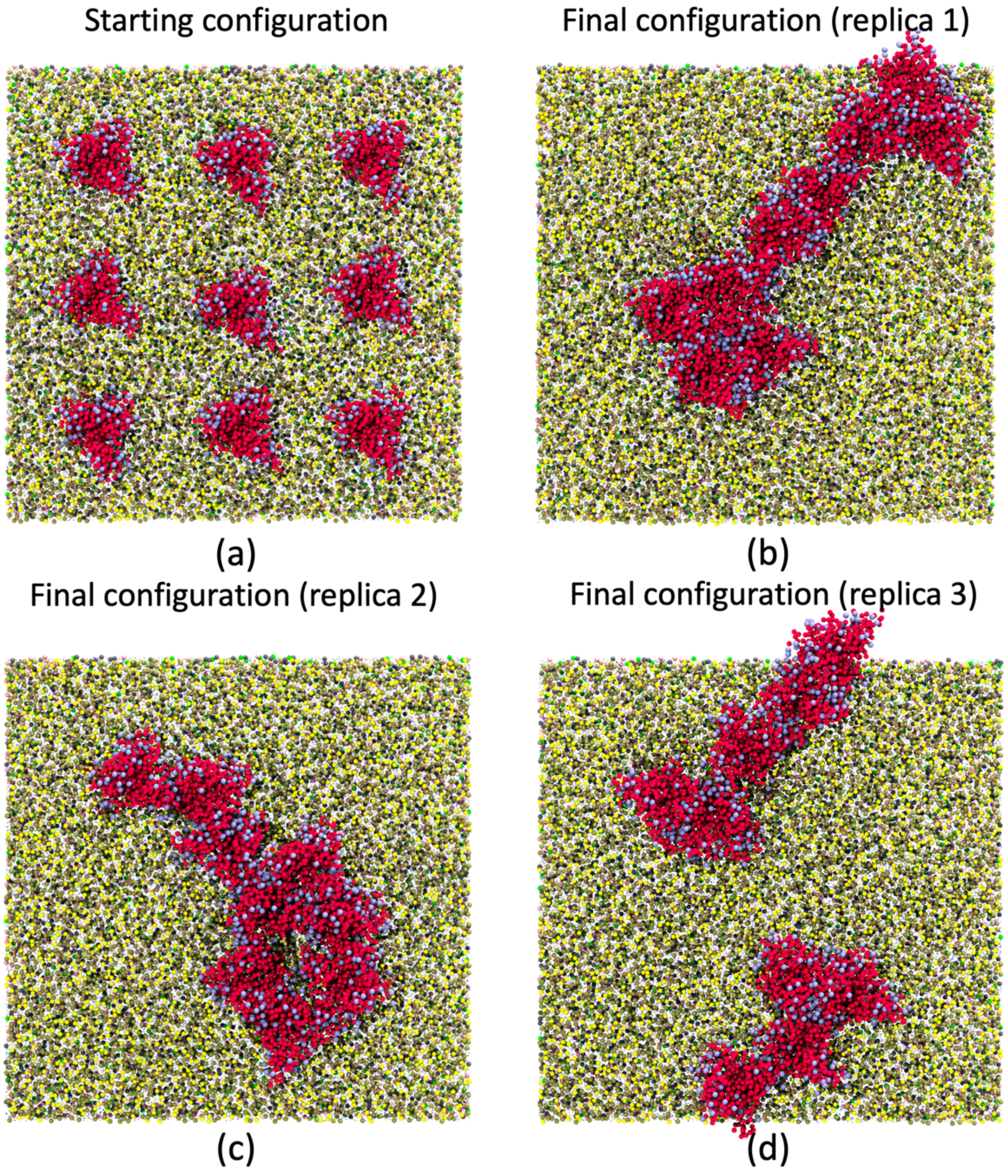


**Figure S8:** (a) Initial configurations and (b, c, d) final configurations from three independent replicas of CG simulations of 9 Env proteins in flat asymmetric bilayers. Env, glycans, POPC, LSM, and cholesterol, POPE, POPS, PIP2 are represented by red, ice blue, yellow, tan, blue, gray, pink, and green, respectively.

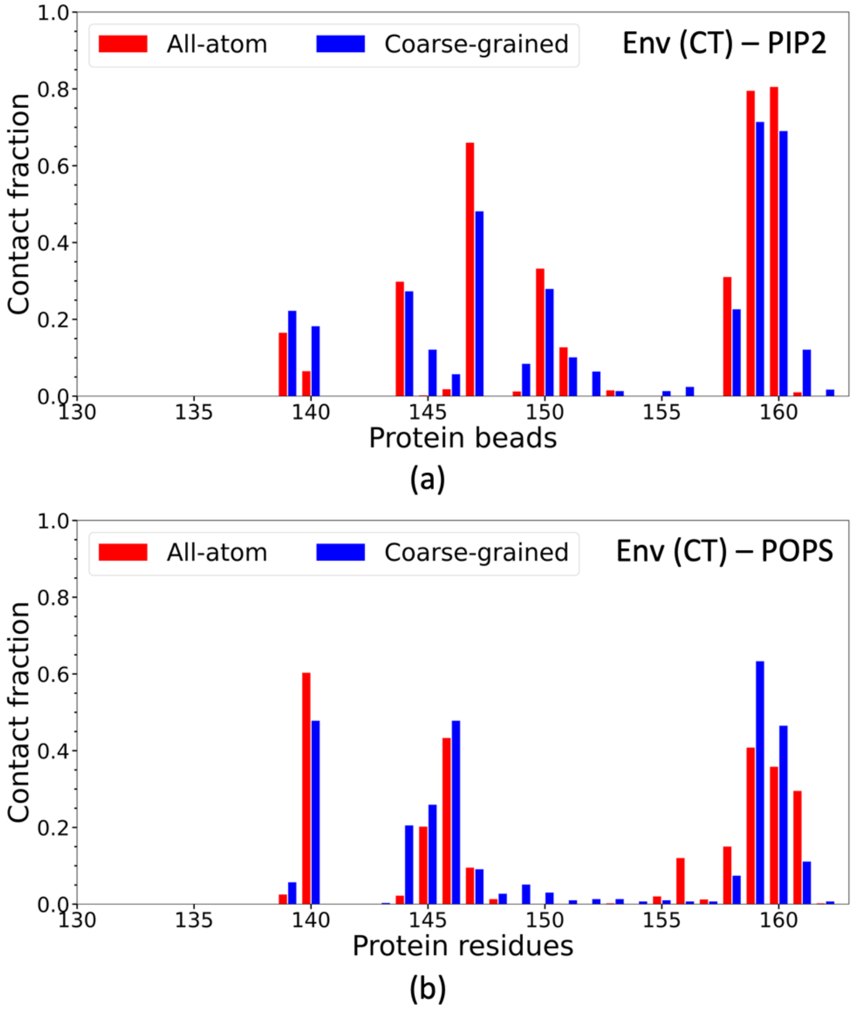


**Figure S9:** Comparison of interactions of (a) PIP2 - CT domain and (b) POPS - CT domain obtained from AA simulation and CG simulations after final REM iteration, where a contact fraction value of 1 indicates a contact maintained between the CG bead and lipids throughout the trajectory.

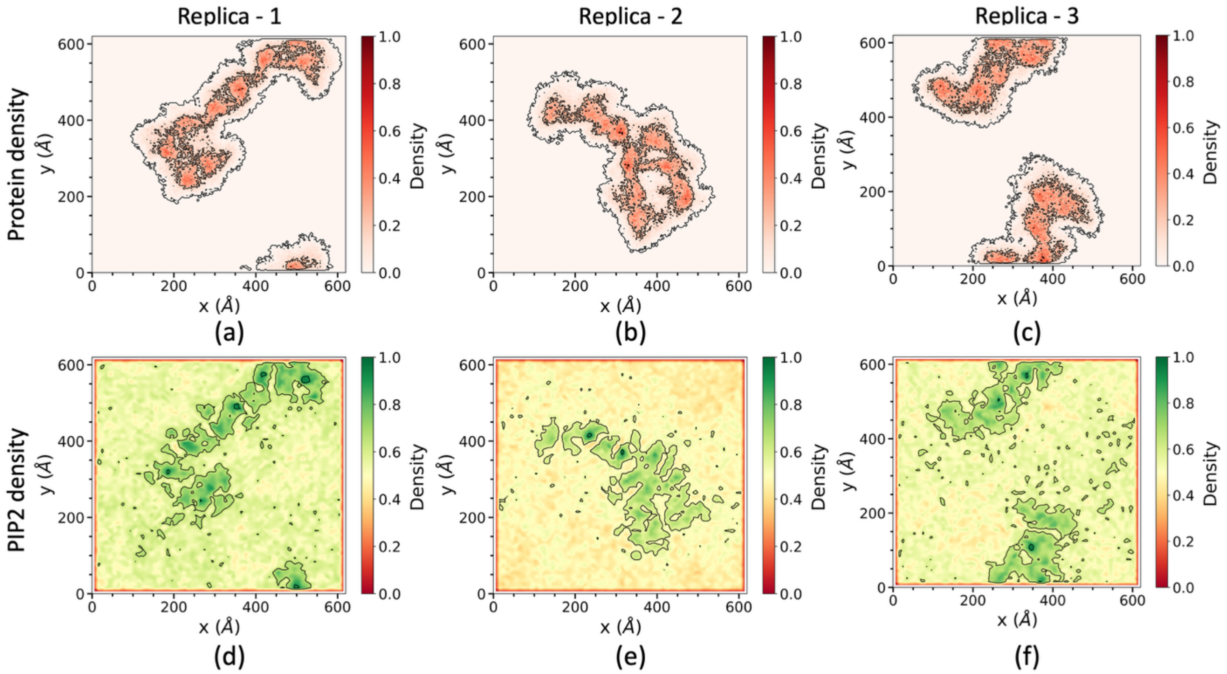


**Figure S10:** Density of Env proteins and PIP2 from three CG independent replica simulations of nine Env proteins in flat asymmetric bilayers. Panels (a-c) show the Env density for replicas 1-3, respectively, and panels (d-f) show the corresponding PIP2 density for replicas 1-3.

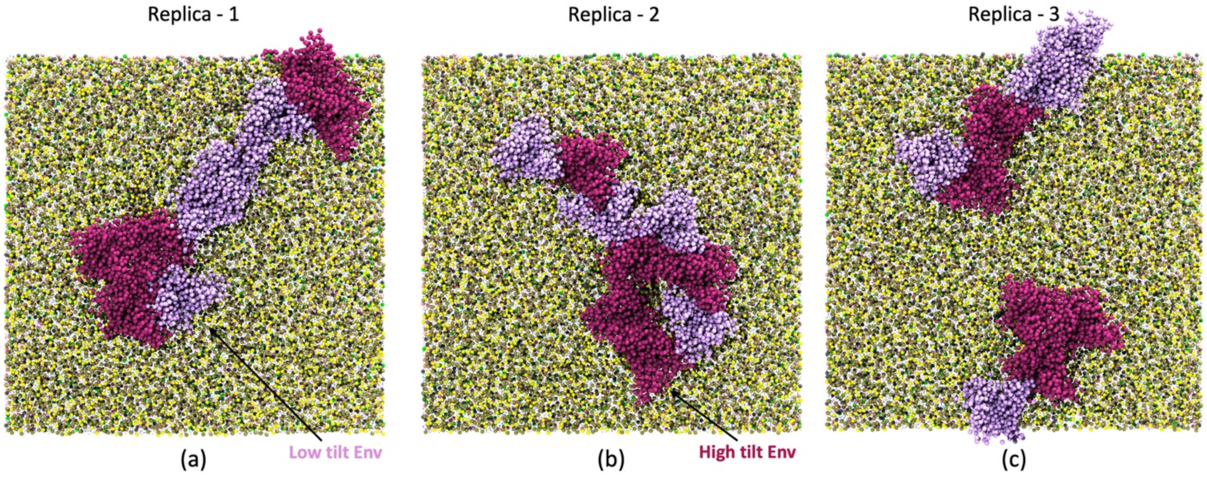


**Figure S11:** Schematic representation of high-tilt (purple) and low-tilt (ice-blue) Env proteins obtained from three independent replicas of coarse-grained simulations of 9 Envs in flat asymmetric bilayers. POPC, LSM, and cholesterol, POPE, POPS, PIP2 are represented by yellow, tan, blue, gray, pink, and green, respectively.

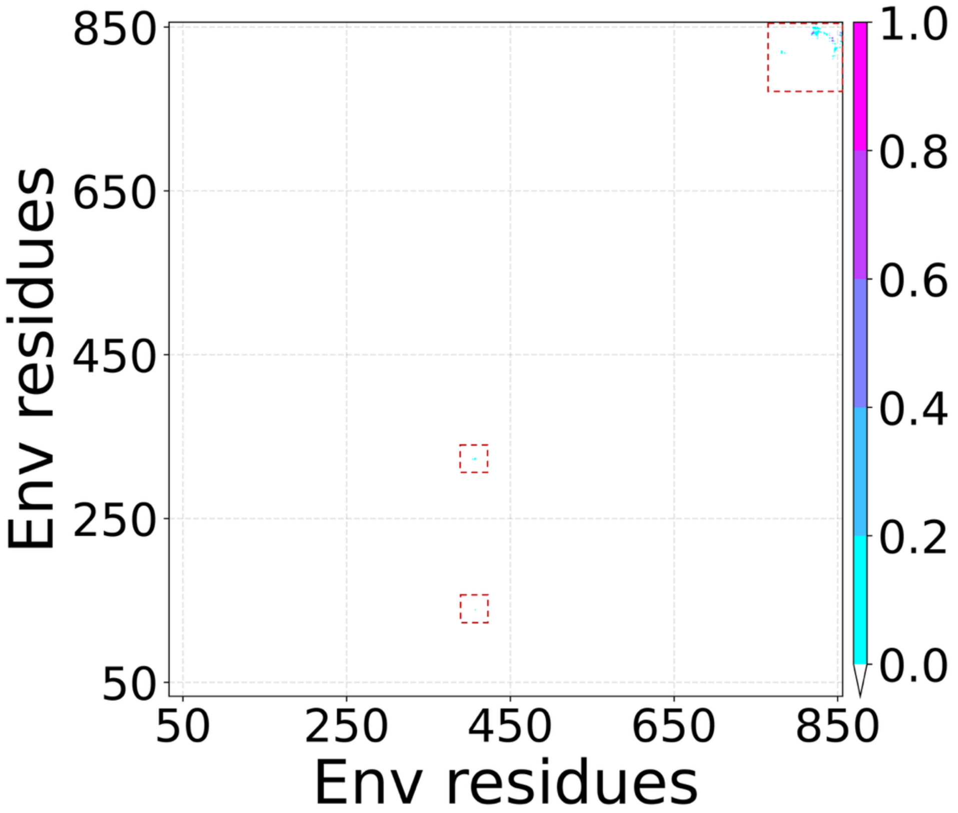


**Figure S12:** Contact map describing interactions between backmapped glycosylated Env dimers from CG simulations, obtained from AA simulations in asymmetric bilayers.
